# SAM/BAM format v1.5 extensions for *de novo* assemblies

**DOI:** 10.1101/020024

**Authors:** Peter J. A. Cock, James Bonfield, Bastien Chevreux, Heng Li

## Abstract

**Summary:** The plain text Sequence Alignment/Map (SAM) file format and its companion binary form (BAM) are a generic alignment format for storing read alignments against reference sequences (and unmapped reads) together with structured meta-data (Li *et al.*, 2009). Driven by the needs of the 1000 Genomes Project which sequenced many individual human genomes, early SAM/BAM usage focused on pairwise alignments of reads to a reference. However, through the CIGAR P operator multiple sequence alignments can also be preserved. Herein we describe clarifications and additions in version 1.5 of the specification to facilitate storing *de novo* sequence alignments: Padded reference sequences (with gap characters), annotation of reads or regions of the reference, and the option of embedding the reference sequence within the file.

**Availability:** The latest public release of the specification is at http://samtools.sourceforge.net/SAM1.pdf, with in development drafts at https://github.com/samtools/hts-specs/ under version control.

## 1 INTRODUCTION

The era of ‘second generation sequencing’ has been one of great change, not just in terms of biological insights and novel algorithms, but also at the practical level of sequence file formats. Here the increasing volumes of high throughput sequencing (HTS) data have pushed early plain text based files formats to their limits, driving the need for efficient alternatives.

In just a few years, SAM/BAM became the *de facto* standard for aligning high throughput sequencing (HTS) reads to reference sequences. It is the default output for tools like BWA (Li and Durbin, 2009) and Bowtie2 (Langmead and Salzberg, 2012), and it is difficult to imagine any new mapping software not offering this as an output format since to do so would handicap downstream analysis and so deter users. This dominance has come about though a combination of factors both social and technical – it would not have been possible without the highly scalable design with BAM random access indexing. This has been greatly helped by two open source reference library implementations, Picard for Java, and the samtools C API (Li *et al.*, 2009). Bindings for the later have been written for major higher level programming languages including Python, Perl, Ruby and R. Additionally the SAM/BAM ecosystem has spurred innovation in visualisation software, with tools like IGV (Robinson *et al.*, 2011) and Tablet (Milne *et al.*, 2013) being popular examples, both written in Java using Picard. As a consequence of this, the best way for any potential successor format such as CRAM (Cochrane *et al.*, 2013) to be adopted will be seamless interoperability with files produced by the samtools and Picard libraries.

For HTS *de novo* assemblies, there has been little agreement to date on file formats beyond simple FASTA for contig sequences, and sometimes FASTQ (Cock *et al.*, 2010) if consensus quality scores are calculated. Partly this reflects the great diversity of *de novo* assembly algorithms, and the resulting variation in the kind and detail of information available. Specifically, de Bruijn graph based assemblers such as Velvet (Zerbino and Birney, 2008) do not normally track individual reads, so it does not make sense to ask for an assembly file showing where each read was placed. One workaround is to perform the assembly and then (re)map the reads to it, giving a SAM/BAM mapping file with useful information about read coverage variation and so on. The idea that SAM/BAM could be extended to store a graph based representation of an assembly was mooted, preserving the linear representation of contigs with reads mapped to them, but annotating how they are interconnected through scaffolding or assembly ambiguities. This has not been pursued as yet. However the participants in the Assemblathon competitions (Earl *et al.*, 2011; Bradnam *et al.*, 2013) have defined a FASTA-like graph format, FASTG (http://fastg.sourceforge.net/). Because this is designed to reduce to a set of linear contigs, perhaps SAM/BAM with reference sequences in FASTG format will become a popular combination?

For more traditional overlap-consensus based assemblers (which may come back into the fore as HTS read lengths are expected to increase), the ACE and CAF file formats have been widely used, for example by PHRAP and CONSED (Gordon *et al.*, 1998), CAP3 (Huang and Madan, 1999), the Roche ‘Newbler’ gsAssember (454 Life Sciences, Roche Applied Science, Branford, CT) and MIRA (Chevreux *et al.*, 1999). These file formats do not scale well with current data volumes, but were still used in part because of features not previously possible in SAM/BAM, which have been addressed with the extensions herein.

### 1.1 Padded reference

The SAM/BAM format has traditionally been used with an unpadded reference sequence (no gap characters), requiring the use of the CIGAR (compact idiosyncratic gapped alignment report) pad (P) operator to capture inter-sequence alignment information implicitly. The Gap5 assembly viewer and editor (Bonfield and Whitwham, 2010) was one of the few tools to implement this. This complexity can be avoided by using a padded reference sequence, where instead CIGAR deletion (D) operators are used to mark where a read does not have the gap found in the reference. This approach has been possible historically, and has now been formally recognised as part of the SAM/BAM standard. Some tools and viewers could already handle this, for instance Tablet (Milne *et al.*, 2013) which supported padded and unpadded reference coordinates as this is inherent in older formats like ACE.

This is illustrated in Table 1 which shows a multiple sequence alignment with the POS and CIGAR entries required in SAM using a padded or unpadded reference. As an example, using the unpadded reference it is important to distinguish the inserts of `C` in Read02 and Read18, or `T` in Read26 as `C----`, `----C` and `--T--` respectively. Using just 1I (one base inserted) in the CIGAR is not sufficient. Similarly for Read11 and Read12, using `4I` does not capture the difference in alignment between `CATA-` and `-ATAC.` All the reads spanning this region should include pad operators consistent with the full 5bp insertion in this example. Note that it is possible to have inconsistent padding, especially if the files have been post-processed or merged. Read08 and Read14 demonstrate correct alignments containing adjacent insertion and deletion CIGAR operators.

**Table 1.**
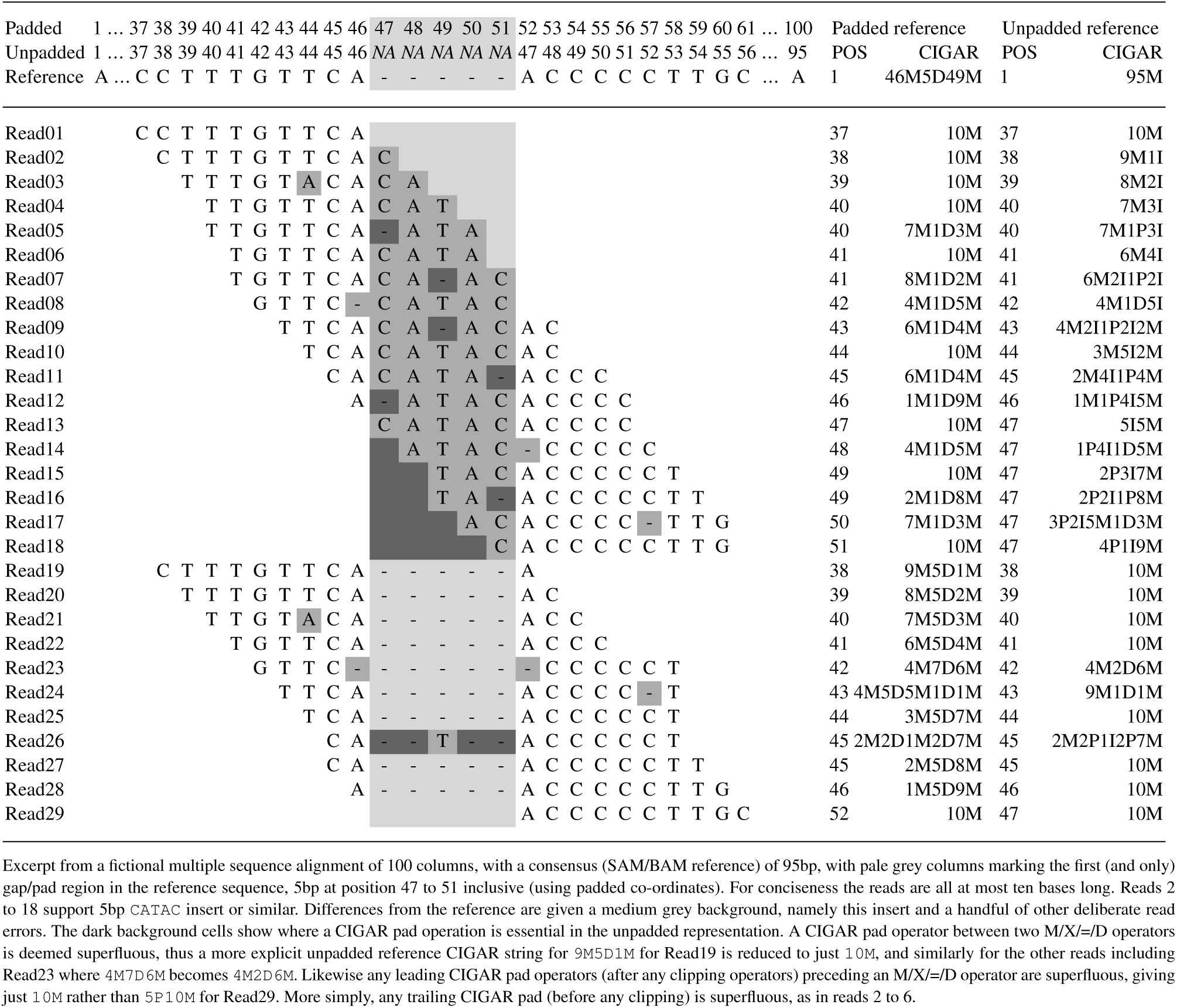
An example gapped alignment

It is expected that updating existing assembly tools to produce SAM/BAM with a padded reference will be simpler than using the unpadded reference, but that the later will continue to be the main focus of downstream analysis. To facilitate this an (experimental) depad command has been added to samtools, together with a wrapper (http://toolshed.g2.bx.psu.edu/view/peterjc/samtools depad) for use within the Galaxy platform (Goecks *et al.*, 2010). This has been tested with the unpadded SAM files produced by the MIRA v4.0 assembler.

Note that the ArchiveBAM format currently strips out the CIGAR P operator, reducing a multiple sequence alignment to a collection of pairwise alignments.

### 1.2 Annotation

One of the strengths of ACE and CAF as assembly file formats is their support for annotation tags intended to guide manual finishing. For example, a potential repeat region identified during assembly can be explicitly marked as such by the assembler. The MIRA assembler (Chevreux *et al.*, 1999) uses a broad collection of such annotation, which is intended to be viewed in a suitable editor like Gap5 (Bonfield and Whitwham, 2010). The SAM/BAM format now defines dummy reads (with no sequence) to hold annotations about a particular region, and additional tags for storing annotations about an individual read. This has been done with a view to extensibility, which should suffice even for rich annotation such as GFF3 format. This may prove useful for reference guided assemblies allowing the assembler to automatically transfer pertinent annotation from the reference genome. Because region specific annotations are held as SAM/BAM reads, they benefit from the existing indexing techniques designed for paired end or strobe reads - including support for features split between contigs (references in SAM/BAM).

### 1.3 Embedded reference

When working with mapping data to a model organism, it makes sense to store the reference sequence outside the SAM/BAM file. However, for *de novo* assemblies, or mapping to draft genomes, using a separate FASTA file complicates data management. In the interests of a single self contained assembly, the SAM/BAM format now defines how to optionally embed each reference (i.e. assembly consensus) sequence as a long dummy read.

## 2 CONCLUSION

These additions should make SAM/BAM a sensible file format choice for HTS assembly tools where individual reads are tracked, particularly more traditional overlap-consensus based assemblers (which may come back into the fore as HTS read lengths continue to increase).

Roche Life Sciences already support unpadded BAM output from some of their ‘Newbler’ 454 sequencing suite, but the extensions to the file format described here would allow a fuller representation. Version 4 of the MIRA assembler supports padded SAM output, using the annotation features. Using samtools these padded SAM assemblies can be converted to unpadded SAM/BAM, and then used with a wide range of downstream analysis tools for tasks such as SNP finding.

Gap5 can now be used to view and edit SAM/BAM assemblies using these new features, and takes full advantage of the annotation present. This is particularly important for manual finishing where the assembler (e.g. MIRA) may tag regions of concern (e.g. identified repeat regions). This replaces existing workflows from MIRA to Gap4 using the CAF file format.

It is our expectation that SAM/BAM can now be used to replace existing first generation sequencing era assembly file formats ACE and CAF, which are inappropriate for current data volumes, and that SAM/BAM will continue to evolve in the future.

## ACKNOWLEDGEMENT

*Funding*: PJAC is funded by the Scottish Government’s Rural and Environment Science and Analytical Services (RESAS) Division.

